# Auranofin and reactive oxygen species inhibit protein synthesis and regulate the level of the PLK1 protein in Ewing sarcoma cells

**DOI:** 10.1101/2024.05.13.593567

**Authors:** Joseph A. Haight, Stacia L. Koppenhafer, Elizabeth L. Geary, David J. Gordon

## Abstract

Novel therapeutic approaches are needed for the treatment of Ewing sarcoma tumors. We previously identified that Ewing sarcoma cell lines are sensitive to drugs that inhibit protein translation. However, translational and therapeutic approaches to inhibit protein synthesis in tumors are limited. In this work, we identified that reactive oxygen species, which are generated by a wide range of chemotherapy and other drugs, inhibit protein synthesis and reduce the level of critical proteins that support tumorigenesis in Ewing sarcoma cells. In particular, we identified that both hydrogen peroxide and auranofin, an inhibitor of thioredoxin reductase and regulator of oxidative stress and reactive oxygen species, activate the repressor of protein translation 4E-BP1 and reduce the levels of the oncogenic proteins RRM2 and PLK1 in Ewing and other sarcoma cell lines. These results provide novel insight into the mechanism of how ROS-inducing drugs target cancer cells via inhibition of protein translation and identify a mechanistic link between ROS and the DNA replication (RRM2) and cell cycle regulatory (PLK1) pathways.

## Introduction

Ewing sarcoma is an aggressive bone and soft-tissue tumor that is characterized by a recurrent chromosomal translocation involving the *EWSR1* and *FLI1* genes (1–3). The current treatment for Ewing sarcoma, which consists of cytotoxic chemotherapy combined with surgery and/or radiation, has changed very little in the past two decades and is associated with suboptimal outcomes (1,4,5). The EWS-FLI1 oncoprotein, which is required for tumorigenesis and only expressed in tumor cells, is an attractive therapeutic target in Ewing sarcoma, but has proven difficult to directly target (6– 8). A complementary approach is to identify unique vulnerabilities incurred by the EWS-FLI1 oncoprotein (9,10).

Recent studies identified protein translation as a potential therapeutic vulnerability and target in Ewing sarcoma tumors (11,12). However, translational and therapeutic approaches to inhibit protein synthesis in tumors are lacking (13). Notably, reactive oxygen species (ROS), which are generated by a range of drugs via different mechanisms, have been linked to inhibition of protein synthesis in cancer cells (14–17). Furthermore, Ewing sarcoma tumors are reported to be sensitive to reactive oxygen species and to the manipulation of oxidative stress (18–24). For example, treatment of Ewing sarcoma tumors with fenretinide, a synthetic retinoid, causes ROS-dependent toxicity in vitro and in vivo in xenograft experiments (21,22). Similarly, auranofin, an inhibitor of thioredoxin reductase and regulator of ROS, was identified using a multipronged screening approach as a drug with activity against Ewing sarcoma tumors (23). Moreover, from a mechanistic standpoint, *SOX6*, a transcriptional target of the EWS-FLI1 oncoprotein, was shown to increase ROS and oxidative stress in Ewing sarcoma cells via upregulation of thioredoxin interacting protein (TXNIP), an inhibitor of thioredoxin and inducer of ROS (24,25). However, despite the sensitivity of Ewing sarcoma cells to ROS-inducing drugs, it is unknown whether ROS targets protein synthesis in this and other sarcoma types.

We began the current study with the aim to test the hypothesis that reactive oxygen species regulate protein synthesis in Ewing sarcoma cells. We found that both hydrogen peroxide and auranofin inhibit protein translation in Ewing and other sarcoma cells. Moreover, from a mechanistic standpoint, we also identified that ROS activates eukaryotic translation initiation factor 4E binding protein 1 (4E-BP1), a repressor of protein translation that binds to the eukaryotic translation initiation factor eIF4E and prevents the formation of the translation initiation complex (26,27). Activation of 4E-BP1 reduces the levels of the oncogenic proteins ribonucleotide reductase M2 (RRM2) and polo-like kinase 1 (PLK1) in Ewing and other sarcoma cell lines. Overall, these results define a link between ROS and protein synthesis in sarcoma cells and identify a mechanistic connection between ROS and the DNA replication (RRM2) and cell cycle regulatory (PLK1) pathways.

## Materials and Methods

### Cell lines and culture

Cell lines were maintained at 37° C in a 5% CO_2_ atmosphere. The A673, TC71, and EW8 cell lines were provided by Dr. Kimberly Stegmaier (Dana-Farber Cancer Institute, Boston, MA), the CB-AGPN cell line was obtained from the Childhood Cancer Repository (Children’s Oncology Group), the HT1080 (RRID:CVCL_0317) and U2OS (RRID:CVCL_0042) cell lines were obtained from ATCC, and the RD cell line was provided by Dr. Munir Tanas (University of Iowa, Iowa City, IA). The non-transformed BJ-tert (RRID:CVCL_6573) and RPE-tert (RRID:CVCL_4388) cell lines were obtained from ATCC. Cells were grown in Dulbecco’s Modified Eagle’s Media (DMEM) supplemented with 10% FBS, 100 IU ml^-1^ penicillin and 100 μg ml^-1^ streptomycin. DNA fingerprinting confirmation of cell lines was performed using the short tandem repeat method and cell lines were used within 8-10 passages of thawing.

### Chemical compounds

Auranofin and TAK-228 (sapanisertib) were obtained from MedChemExpress. N-acetylcysteine, hydrogen peroxide, and doxycycline were obtained from Sigma. Puromycin was obtained from ThermoFisher Scientific. Auranofin was dissolved in DMSO and then used at a 1:1000 dilution in assays. TAK-228 was also prepared in DMSO and used at a 1:1000 dilution in assays. N-acetylcysteine was prepared fresh for each experiment in DMEM media.

### Cell viability and drug dose response assays

Cell proliferation was measured using the AlamarBlue (resazurin; Sigma) fluorescence assay, as previously described (11,12,28,29). Briefly, approximately 5 x 10^4^ cells were plated per well of a 96-well plate. Cells were then treated the next day with a range of drug concentrations for 72 hours. Fluorescence measurements were obtained, after adding the AlamarBlue reagent, using a FLUOstar Omega microplate reader (BMG Labtech). IC50 values were calculated using log-transformed and normalized data (GraphPad Prism 10.1.0). Cell viability was quantified using 0.4% Trypan Blue (ThermoFisher), which stains dead cells, with a DeNovix Cell Drop BF automated cell counter.

### Puromycin labeling

Protein synthesis was assessed using puromycin labeling (SUnSET technique), as described (11,12,30,31). For labeling of newly synthesized proteins, puromycin (2 μg/mL) was added to cells at a 1:400 dilution. The cells were then incubated with the puromycin for one hour before cell lysates were collected, as described in the Immunoblotting section. Protein loading for the immunoblots was normalized using cell number.

### O-propargyl-puromycin labeling

Protein synthesis was assessed using O-propargyl-puromycin Click-iT labeling according to the manufacturer’s instructions (ThermoFisher Scientific) (32,33). Flow cytometry was performed using a Becton Dickinson LSR II instrument. Briefly, cells were incubated with O-propargyl-puromycin (1:1000, 10 μM final concentration) for one hour. Cells were then collected using trypsin and fixed in a 4% paraformaldehyde fixative solution. Cells were washed by centrifugation and resuspended in Click-iT saponin-based permeabilization and wash reagent. Click-iT Plus reaction cocktail was added to the cell suspension for thirty minutes, after which the cells were washed by centrifugation. Flow cytometry was performed on a Becton Dickinson LSR II flow cytometer.

### Protein isolation and immunoblotting

Protein extracts for immunoblotting were prepared by incubating cells in RIPA buffer (Boston BioProducts), supplemented with protease and phosphatase inhibitors (Halt Protease & Phosphatase Inhibitor Cocktail, EDTA-free; ThermoFisher Scientific), for 20 minutes. Supernatants were collected following a 15-minute centrifugation at 17,000 r.c.f. at 4°C. SDS-PAGE was used to separate proteins, which were then transferred to polyvinylidene difluoride membranes (Millipore). Antibodies to the following proteins were used in the immunoblots: puromycin (Millipore, #AF488, 1:2000; RRID:AB_2737590), 4E-BP1 (Cell Signaling, #9644, 1:1000; RRID:AB_2097841), p-4E-BP1-37/46 (Cell Signaling, #2855, 1:1000; RRID:AB_560835), RRM2 (Santa Cruz Biotechnology, #398294, 1:500; RRID:AB_2894824), PLK1 (Cell Signaling, #4513, 1:1000; RRID:AB_2167409), tubulin (Proteintech, #66031, 1:2000; RRID:AB_28834830), and vinculin (Proteintech, #26520, 1:5000; RRID:AB_2919877). Protein loading for the immunoblots was normalized using cell number.

### Reactive oxygen species detection

ROS was quantified using a DCFDA (2’,7’-dichlorofluorescin diacetate) assay kit according to the manufacturer’s instructions (Abcam). Briefly, 1 x 10^4^ cells were plated per well of a 96-well plate in phenol red-free media. Drugs were then added the next day and incubated for the amount of time indicated in the figure legends. The DCDFA reagent, which was dissolved in the kit buffer at a 20 mM concentration, was added to the wells at a 1:800 dilution and then incubated for 45 minutes at 37° C. Fluorescence measurements (excitation 485 nM/emission 535 nM) were obtained using a FLUOstar Omega microplate reader (BMG Labtech).

### EdU labeling

Detection of DNA replication was performed in duplicate using a Click-iT EdU-488 kit (ThermoFisher Scientific) (34). Briefly, cells were labeled with 10 μM EdU for one hour and then harvested using trypsin and fixed using the Click-iT fixative buffer. Cells were then washed by centrifugation and resuspended in Click-iT saponin-based permeabilization and wash reagent. Click-iT Plus reaction cocktail was added to the cell suspension for thirty minutes, after which the cells were washed by centrifugation. Flow cytometry was performed on a Becton Dickinson LSR II flow cytometer.

### Doxycycline-inducible 4E-BP1-Ala

The full-length 4E-BP1 cDNA, with alanine substitutions at Thr37, Thr46, Ser65, and Thr70 and a FLAG-tag, was obtained as a gene block (IDT; Coralville, IA) and cloned into a lentiviral vector as described in previous publications (12,34).

### Reverse phase protein array (RPPA)

RPPA analysis of cell lines were performed by the RPPA Core Facility at the MD Anderson Cancer Center. Cells were provided to the core facility as frozen pellets and the protein extraction, data normalization, and analysis were performed according to facility protocols (https://www.mdanderson.org/research/research-resources/core-facilities/functional-proteomics-rppa-core/education-and-references.html) (RRID:SCR_016649).

### Statistical Analysis

Student’s t-test two-tailed was used to calculate P-values for the comparison of two groups. Statistical analyses were conducted using GraphPad Prism 10.1.0 (RRID: SCR_002798).

## Results

### Hydrogen peroxide inhibits protein synthesis in sarcoma cell lines

To directly test the hypothesis that ROS inhibits protein synthesis in Ewing sarcoma cells we treated two cell lines, EW8 and TC71, with hydrogen peroxide for three hours and then measured nascent protein synthesis using puromycin labeling (SUnSET technique) and immunoblotting (30,31). Figure 1A shows that hydrogen peroxide decreases protein synthesis in Ewing sarcoma cells and that this effect is attenuated by co-treatment of the cells with the ROS-scavenger n-acetylcysteine (NAC). We also used a flow cytometry approach (O-propargyl-puromycin) to quantify, in single cells, the effect of hydrogen peroxide on protein synthesis (32,33). Notably, hydrogen peroxide reduced protein synthesis to the same extent as cycloheximide, a well-established inhibitor of protein synthesis that blocks translation elongation, and this effect was blocked by co-treatment with NAC (Figure 1B-C). We then tested additional sarcoma cell lines, HT1080 (fibrosarcoma), RD (rhabdomyosarcoma), and U2OS (osteosarcoma), that represent other sarcoma subtypes and found that hydrogen peroxide also inhibited protein synthesis, similar to the results obtained with the Ewing sarcoma cell lines (Figure 1D).

**Figure 1.**
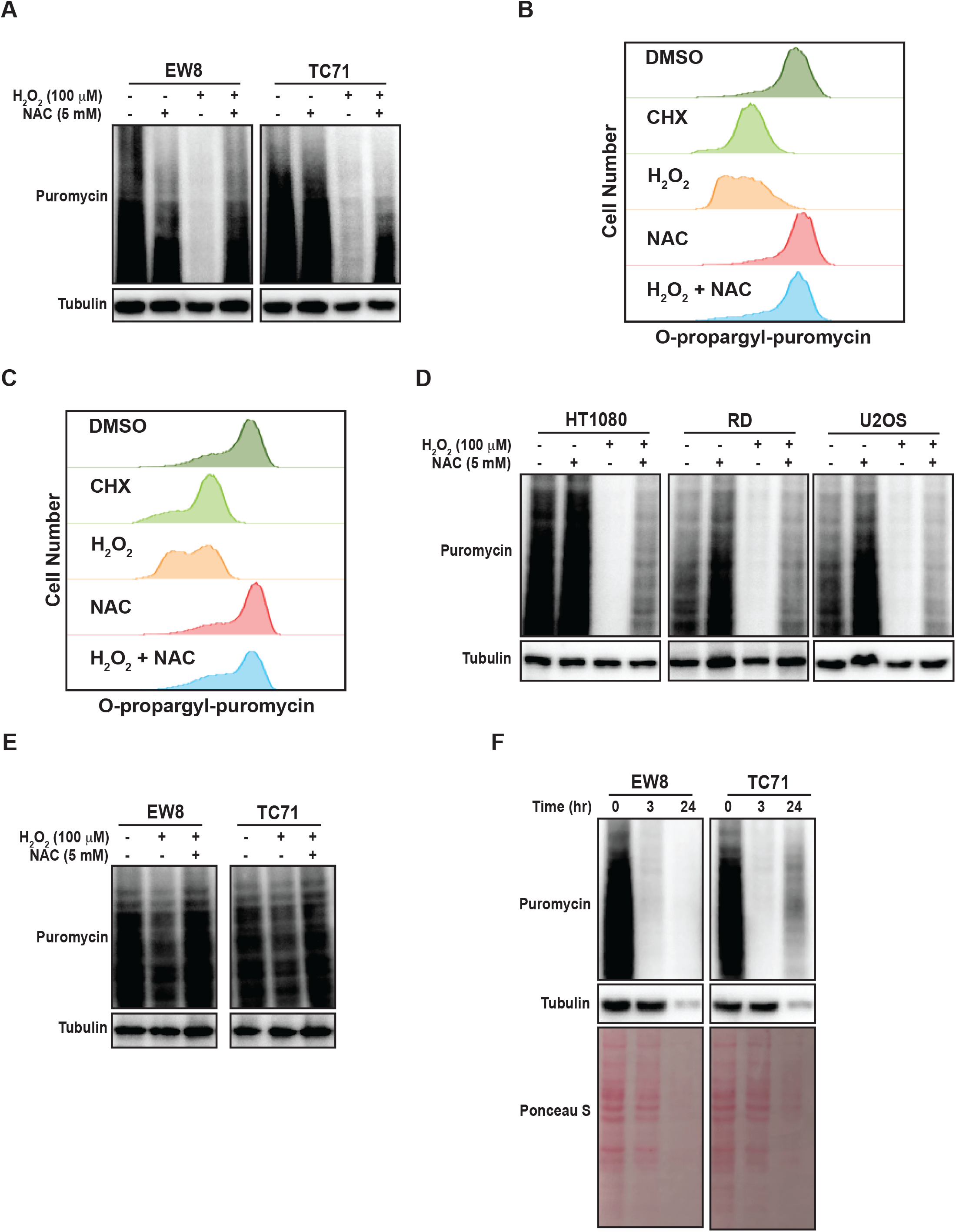
Hydrogen peroxide inhibits protein synthesis in sarcoma cell lines. (A) EW8 and TC71 cells were treated with hydrogen peroxide, NAC, or the combination for three hours. Cells were labeled with puromycin to quantify protein synthesis and then lysates were collected for immunoblotting. (B-C) EW8 (B) and TC71 (C) cells were treated with hydrogen peroxide (100 μM), NAC (5 mM), or the combination for three hours and then labeled with O-propargyl-puromycin for quantification of protein synthesis by flow cytometry. (D) Additional sarcoma cell lines were treated with hydrogen peroxide, NAC, or the combination for three hours and then labeled with puromycin for immunoblotting. (E) EW8 and TC71 cells were treated with hydrogen peroxide for one hour and then protein synthesis was assessed using puromycin. (F) Ewing sarcoma cells were treated with hydrogen peroxide (100 μM) for three hours and then allowed to recover for twenty-one hours before protein synthesis was assessed using puromycin.

Next, we treated the Ewing sarcoma cells lines with hydrogen peroxide for one hour, in contrast to the three hours used in the previous experiments, and identified a significant decrease in protein synthesis at this earlier time point, albeit to a lesser extent than at the later time point (Figure 1E). We also treated Ewing sarcoma cells lines with hydrogen peroxide for three hours and then removed the hydrogen peroxide and allowed the cells to recover for twenty-one hours before measuring nascent protein synthesis. Figure 1F shows that a three-hour treatment with hydrogen peroxide is sufficient to cause prolonged inhibition of protein synthesis, that extends after the hydrogen peroxide is removed. In addition, this more prolonged inhibition of protein synthesis also reduced the level of the protein loading control, tubulin. Furthermore, staining total protein with Ponceau S showed a significant reduction in total protein levels at the later time point. Of note, the loading for these and subsequent immunoblots were normalized using cell number and not total protein due to the effects of the drugs on protein synthesis.

### Auranofin inhibits protein synthesis in Ewing sarcoma cell lines

Auranofin is an inhibitor of thioredoxin reductase and regulator of oxidative stress that is used for the treatment of rheumatoid arthritis (35,36). Auranofin was previously identified, using a multipronged drug screening approach, as a drug that inhibits the growth of Ewing sarcoma cells in vitro and in vivo (23). Of note, this sensitivity of Ewing sarcoma cell lines to auranofin was dependent on expression of the EWS-FLI1 oncoprotein (23). Consistent with this result, analysis of Cancer Dependency Map data (Broad Institute) identifies that Ewing sarcoma cell lines are significantly more sensitive to CRISPR-mediate knockout of thioredoxin reductase than other cancer cell lines (Figure 2A) (37,38). Figure 2B demonstrates that treatment of Ewing sarcoma cells with auranofin increases reactive oxygen species, as detected using the fluorescent ROS-detector DCFDA (2’,7’-dichlorofluorescin diacetate). Auranofin, as previously reported, also decreases the growth of Ewing sarcoma cell lines in a 72 hour growth assay (Figure 2C) (23). Next, we assessed the effect of auranofin dose on protein synthesis using a 24 hour drug treatment and identified that inhibition of protein synthesis occurred in the low (2-5) micromolar dose range (Supplementary Figure 1). To confirm that the effect of auranofin on protein synthesis is mediated by ROS we performed a co-treatment experiment with NAC and found that the ROS scavenger was able to block the effects of auranofin on nascent protein synthesis (Figure 2D). We also used flow cytometry (O-propargyl-puromycin) to investigate the effect of auranofin on protein synthesis in single cells and confirmed that the drug reduces protein synthesis (Figure 2E-F). The effect of auranofin on protein synthesis was then assessed at an earlier time point, 6 hours after drug addition, compared to the timepoint of 24 hours used in the previous experiments. Figure 2G demonstrates that auranofin inhibits protein synthesis in Ewing sarcoma cells as early as 6 hours after drug addition, although the effect is larger with a longer duration of drug treatment. We also used flow cytometry and O-propargyl-puromycin labeling to validate that auranofin inhibits protein synthesis at this earlier (6 hour) time point (Supplementary Figure 1B-C). Cell viability was not significantly compromised at either the 6 hour or 24 hour timepoints (Supplementary Figure 1D-E). We then tested additional sarcoma cell lines that represent other sarcoma subtypes and found that auranofin also inhibited protein synthesis, similar to the results obtained with the Ewing sarcoma cell lines (Figure 2H). Finally, we treated two non-transformed cell lines, BJ-tert and RPE-tert, with auranofin and then labeled newly synthesized proteins with puromycin. Supplementary Figure 2A shows that auranofin inhibits protein synthesis in these non-transformed cell lines, and the effect is rescued by treatment with NAC. However, in contrast to the Ewing sarcoma cell lines, auranofin had minimal effect on the growth of the BJ-tert and RPE-tert cell lines in dose response assays, suggesting that the differential toxicity of the drug with the Ewing sarcoma and non-transformed cells is due to an increased reliance on active protein synthesis in the cancer cells (Supplementary Figure 2B-C).

**Figure 2.**
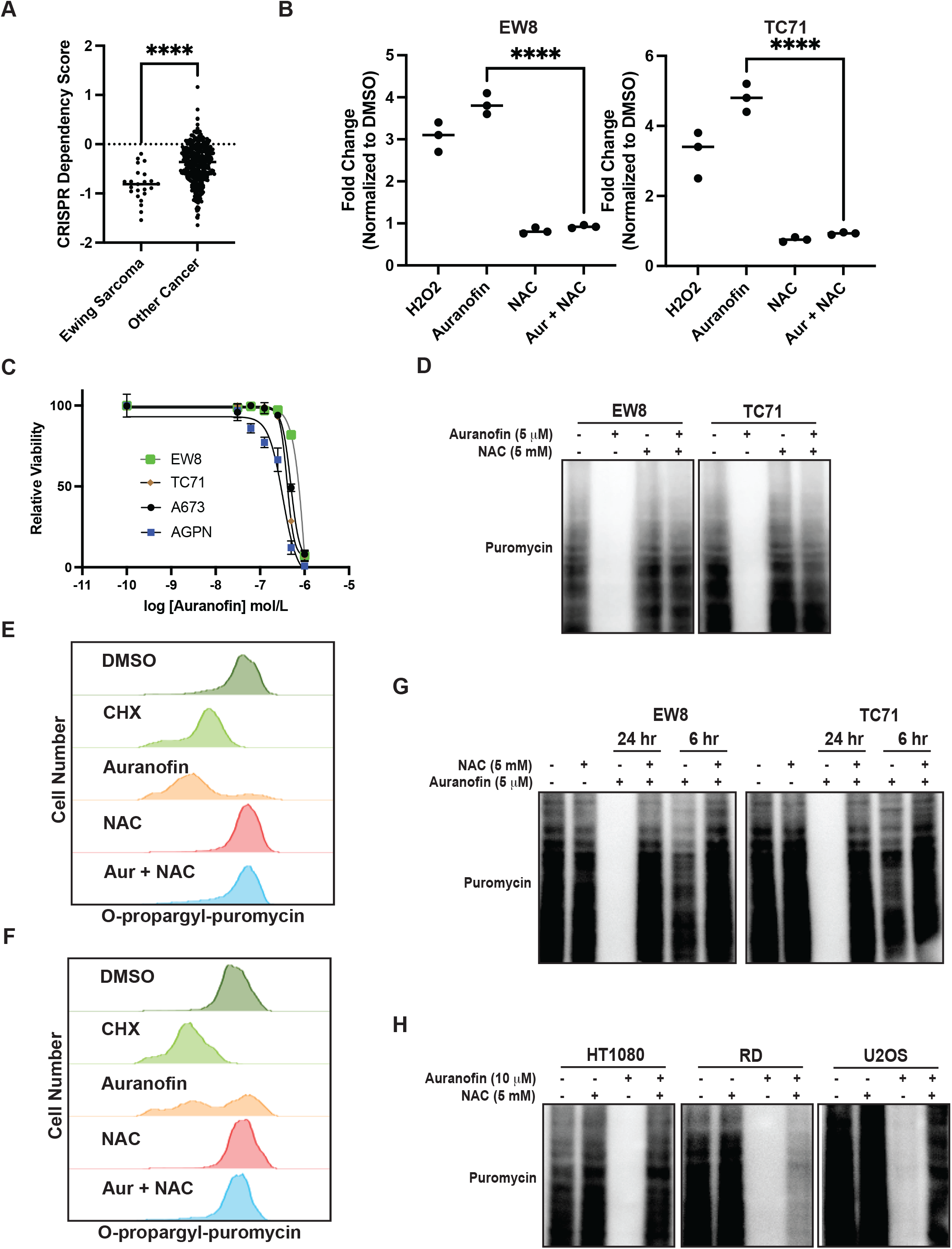
Auranofin inhibits protein synthesis in Ewing sarcoma cell lines. (A) Dependency Map data (https://depmap.org/portal/) showing the effect of CRISPR knockout of thioredoxin reductase on the growth of cancer cell lines. (B) EW8 and TC71 cells were treated with auranofin (5 μM) for twenty-four hours, or hydrogen peroxide (100 μM) for three hours, and then ROS was quantified using a DCFDA fluorescence assay and normalized to cells treated with DMSO. Error bars represent the mean ± SD of three independent experiments with six technical replicates per drug treatment. (C) Dose response curves for Ewing sarcoma cells treated with auranofin. Cell viability was assessed 72 hours after drug was added using the AlamarBlue assay. Error bars represent the mean ± SD of three technical replicates. The results are representative of two independent experiments. (D) EW8 and TC71 cells were treated with auranofin, NAC, or the combination for twenty-four hours. Cells were labeled with puromycin to quantify protein synthesis and then lysates were collected for immunoblotting. (E-F) EW8 (E) and TC71 (F) cells were treated with auranofin (5 μM) for twenty-four hours and then labeled with O-propargyl-puromycin for quantification of protein synthesis by flow cytometry. (G) EW8 and TC71 cells were treated with auranofin, NAC, or the combination for either six or twenty-four hours and then labeled with puromycin to quantify protein synthesis. (H) Additional sarcoma cell lines were treated with auranofin, NAC, or the combination for twenty-four hours and then labeled with puromycin for immunoblotting. Protein loading for the immunoblots was normalized using cell number.

### Auranofin reduces the level of the RRM2 protein and blocks DNA replication Ewing sarcoma cells

In previous work, we identified that Ewing sarcoma cell lines require active protein synthesis to maintain a stable level of the ribonucleotide reductase M2 (RRM2) protein, which is a potential therapeutic target in Ewing sarcoma tumors (12,34,39,40). Consequently, we next evaluated whether auranofin and ROS downregulate RRM2 levels in Ewing sarcoma cells. Figure 3A demonstrates that auranofin depletes RRM2 in Ewing sarcoma cell lines and that this effect is blocked by co-treatment of the cells with the ROS-scavenger NAC. Knockdown or inhibition of RRM2 in Ewing sarcoma is lethal and Figure 3B shows that auranofin significantly impairs the growth of four different Ewing sarcoma cell lines and that this effect is abrogated by co-treatment with NAC. RRM2 is a subunit of ribonucleotide reductase, which is the rate limiting complex in the synthesis of deoxyribonucleotides, and required for progression through S-phase (41–43). Cell cycle analysis demonstrates that auranofin blocks DNA replication, as assessed using EdU labeling, and arrests cells throughout S-phase, as expected for a drug that inhibits ribonucleotide reductase activity (Figure 3C).

**Figure 3.**
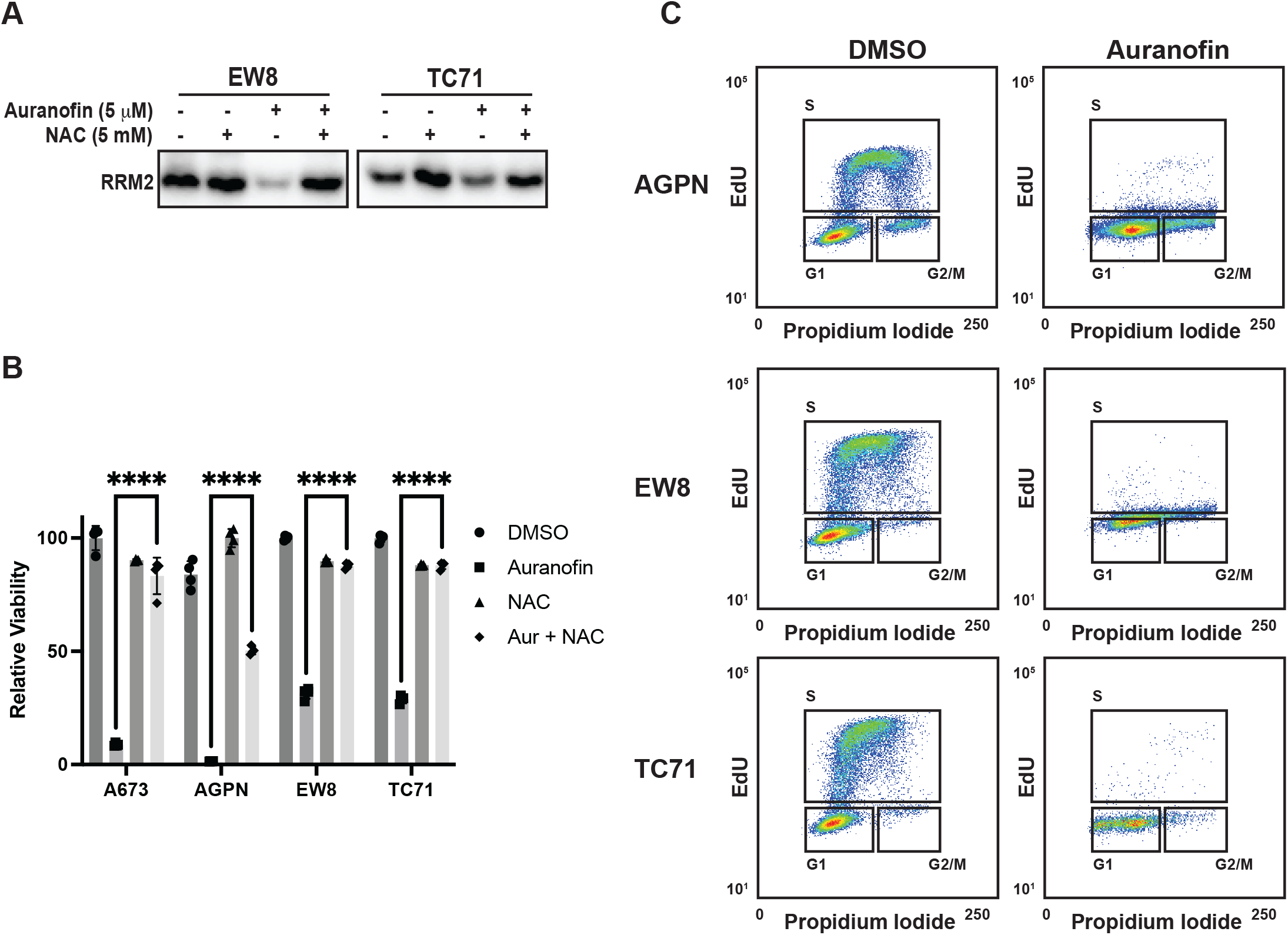
Auranofin reduces the level of the RRM2 protein and blocks DNA replication Ewing sarcoma cells. (A) EW8 and TC71 cells were treated with auranofin, NAC, or the combination for twenty-four hours and then cell lysate was collected for immunoblotting for RRM2. Protein loading for the immunoblots was normalized using cell number. (B) Ewing sarcoma cell lines were treated with auranofin (5 μM), NAC (5 mM), or the combination for 72 hours and then cell viability was quantified using AlamarBlue. Error bars represent the mean ± SD of four technical replicates. The results are representative of two independent experiments. (C) Ewing sarcoma cell lines were treated with auranofin for twenty-four hours and then the cell cycle distribution was analyzed using EdU labeling and propidium iodide.

### Auranofin and reactive oxygen species activate the translational repressor 4E-BP1 in sarcoma cell lines

Reactive oxygen species are reported to inhibit protein synthesis via multiple mechanisms, including modification of cysteines in ribosomal proteins and activation of the translational repressor 4E-BP1 (14–16). Specifically, 4E-BP1 binds to the eukaryotic translation initiation factor eIF4E and prevents the formation of the translation initiation complex (26,27). Notably, in previous work, we identified a role for 4E-BP1, which is regulated by phosphorylation, in blocking protein synthesis in Ewing sarcoma cells (11,12). Consequently, we tested whether ROS activates 4E-BP1 in Ewing sarcoma cells. Figure 4A shows that hydrogen peroxide decreases the phosphorylation of 4E-BP1, which reflects activation of the translational repressor, in Ewing sarcoma cell lines and this effect is blocked by co-treatment with NAC. Similarly, auranofin also inhibits the phosphorylation of 4E-BP1 and, thereby, activates the repressor of cap-dependent protein translation (Figure 4B). To investigate the effect of active 4E-BP1 we previously generated multiple sarcoma cell lines that express a doxycycline-inducible, constitutively-active 4E-BP1 protein with alanine substitutions at the phosphorylation sites Thr37, Thr46, Ser65, and Thr70 (TO-4E-BP1-Ala) (12,44). Figure 4C shows that the expression of constitutively-active 4E-BP1-Ala in Ewing sarcoma cells inhibits protein synthesis and reduces the level of the RRM2 protein, which we previously showed is regulated by 4E-BP1 (12). To identify additional proteins that are regulated by 4E-BP1 we evaluated the effect of the inducible, constitutively-active 4E-BP1-Ala on the proteome of multiple sarcoma cell lines, described in an earlier publication, using reverse phase protein arrays (RPPA) (Figure 4D and Supplementary Table 1) (12). Overlap of the RPPA data for the five cell lines identified that the expression of active 4E-BP1-Ala in all of the sarcoma cell lines results in the downregulation of the level of the PLK1 protein (Figure 4E and Supplementary Table 2).

**Figure 4.**
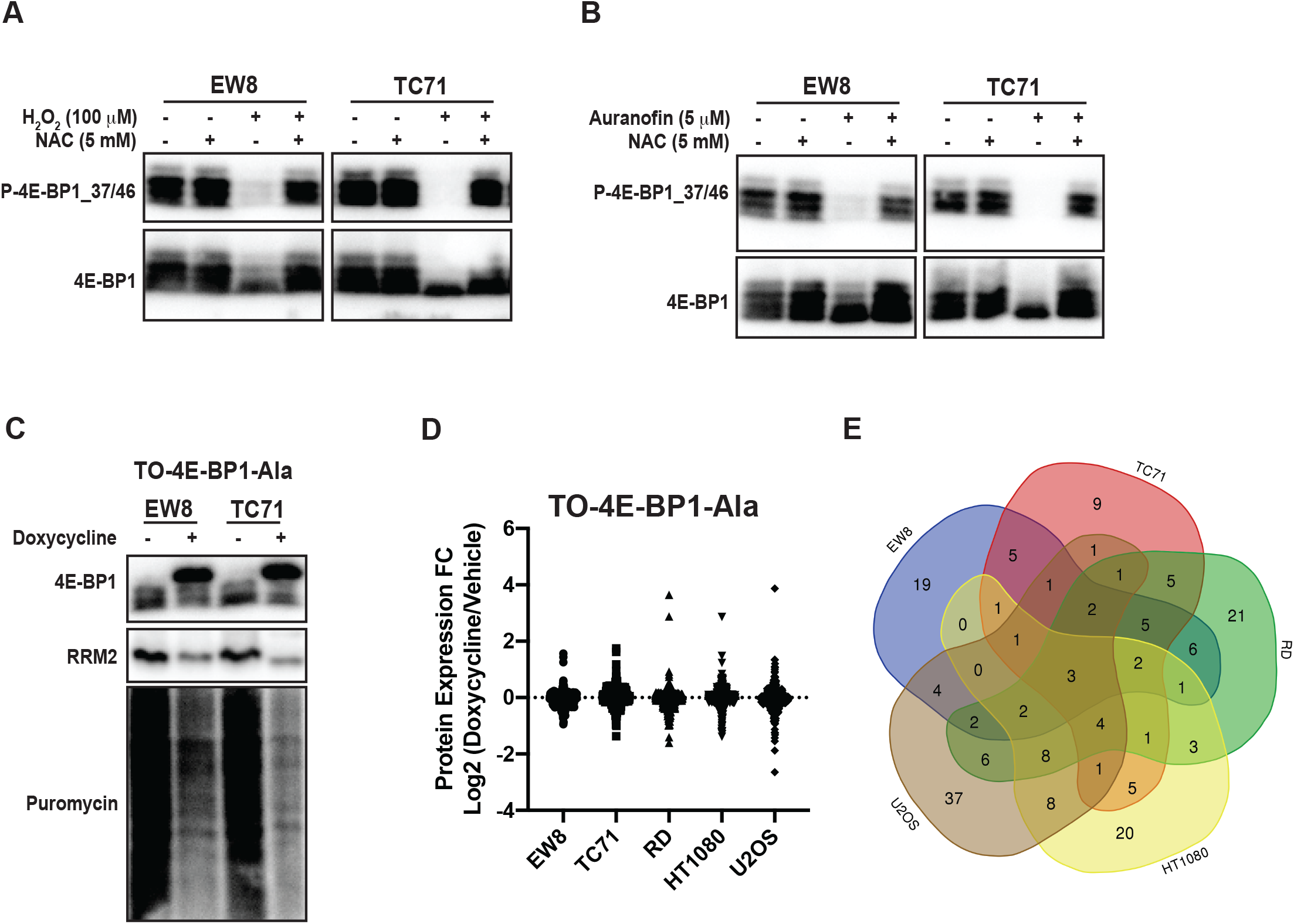
Auranofin and reactive oxygen species activate the translational repressor 4E-BP1 in sarcoma cell lines. (A) EW8 and TC71 cells were treated with hydrogen peroxide, NAC, or the combination for three hours and then cell lysates were collected for immunoblotting. Protein loading for the immunoblots was normalized using cell number. (B) EW8 and TC71 cells were treated with auranofin, NAC, or the combination for twenty-four hours and then cell lysates were collected for immunoblotting. Protein loading for the immunoblots was normalized using cell number. (C) TO-4E-BP1-Ala cell lines were treated with doxycycline for 48 hours to induce expression of 4E-BP1-Ala and then cell lysates were collected after labeling the cells with puromycin. (D) Comparison of protein expression (RPPA) in TO-4E-BP1-Ala sarcoma cell lines treated with vehicle or doxycycline for 48 hours. (E) Venn diagram showing the overlap of the RPPA data for proteins that are downregulated by >1.25-fold in the TO-4E-BP1-Ala sarcoma cell lines in the presence of doxycycline.

### Auranofin and active 4E-BP1 reduce the level of the PLK1 protein in sarcoma cell lines

PLK1 is a well-described therapeutic target in multiple types of cancer, including Ewing sarcoma (45–49). To validate PLK1 as a target of 4E-BP1 we treated the 4E-BP1-Ala cells with doxycycline and performed immunoblotting for PLK1. Figure 5A shows that the expression constitutively-active 4E-BP1 reduces the level of the PLK1 protein, consistent with the RPPA results. Auranofin also reduced the level of the PLK1 protein in Ewing (Figure 5B) and other (Figure 5C) sarcoma cell lines. In previous work, we identified that inhibition of mTORC1/2 activates 4E-BP1 and inhibits protein synthesis in sarcoma cell lines (12). Figure 5D shows that treatment of sarcoma cell lines with the mTORC1/2 inhibitor TAK-228 activates 4E-BP1 and downregulates the level of the PLK1 protein (Figure 5D).

**Figure 5.**
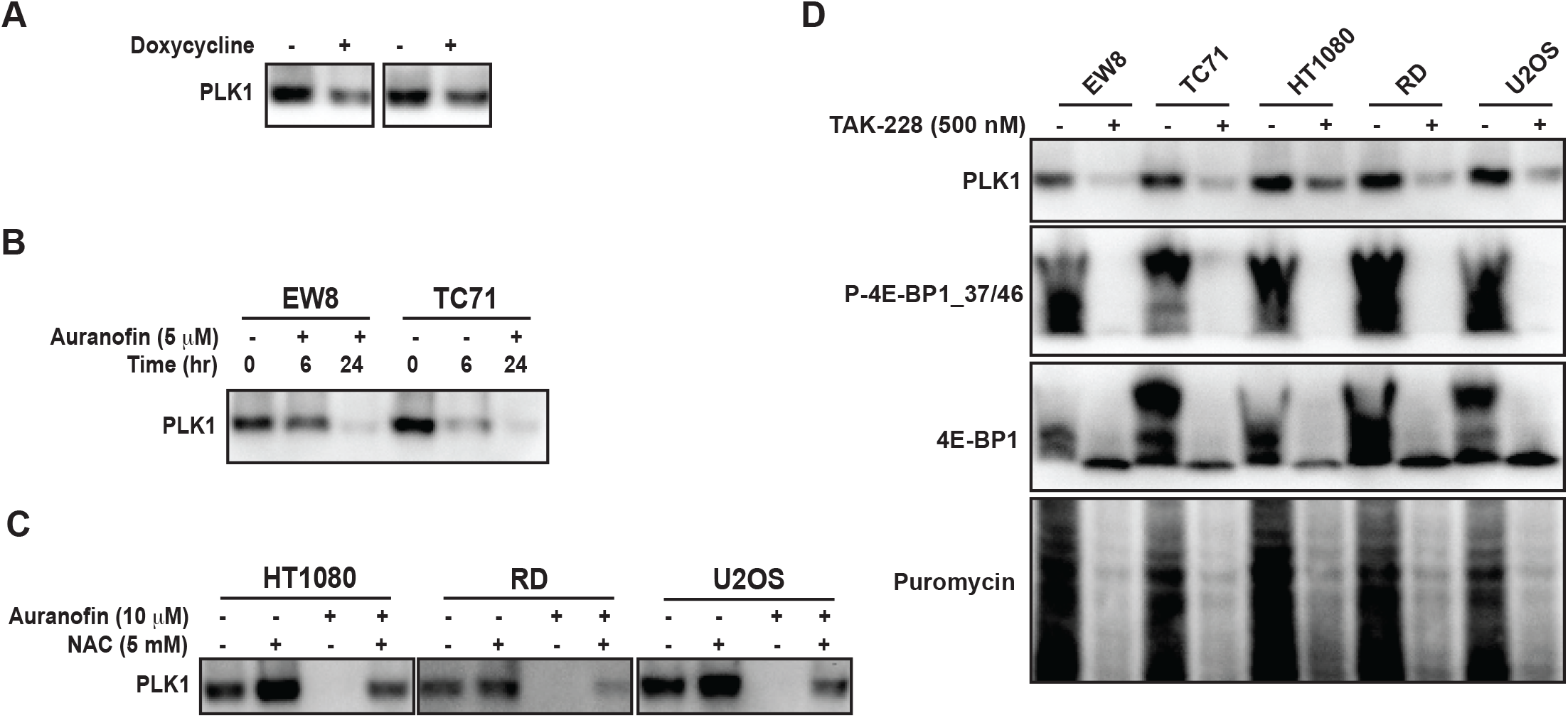
Auranofin and active 4E-BP1 reduce the level of the PLK1 protein in sarcoma cell lines. (A) TO-4E-BP1-Ala cell lines were treated with doxycycline for 48 hours to induce expression of 4E-BP1-Ala and then cell lysates were collected for immunoblotting. (B) EW8 and TC71 cells were treated with auranofin for either six or twenty-four hours and then cell lysates were collected for immunoblotting. (C) Additional sarcoma cell lines were treated with auranofin, NAC, or the combination for twenty-four hours and then cell lysates were collected for immunoblotting for PLK1. (D) Sarcoma cell lines were treated with the mTORC1/2 inhibitor TAK-228 for twenty-four hours and then cell lysates were collected for immunoblotting. Protein loading for the immunoblots was normalized using cell number.

## Discussion

Ewing sarcoma is an aggressive sarcoma of the bone and soft tissue that is treated with chemotherapy in combination with surgery and/or radiation (1,2). Unfortunately, the standard treatment regimen for Ewing sarcoma is unchanged in over two decades and this therapy is associated with significant on- and off-treatment morbidities (4,5). Consequently, there is an urgent need to identify new therapeutic approaches and targets in Ewing sarcoma tumors (8). This study investigated the effects of reactive oxygen species (ROS) on protein synthesis in Ewing sarcoma cell lines. We found that hydrogen peroxide or auranofin, an inhibitor of thioredoxin reductase and ROS inducer, inhibited protein synthesis, activated the repressor of cap-dependent protein translation 4E-BP1, and reduced the levels of the proteins RRM2 and PLK1, both of which promote tumorigenesis in Ewing sarcoma tumors (12,34,39,40,45–47).

Our findings provide novel insight into the mechanism of how ROS-inducing drugs target cancer cells. It is well-established that ROS can damage DNA, leading to apoptosis or cell cycle arrest (50,51). However, our results demonstrate that ROS can also target protein synthesis, which is another essential process for cell survival and proliferation (13). Of note, we found that auranofin also inhibits protein synthesis in non-transformed cells (BJ-tert and RPE-tert), but the drug had minimal effects on the growth and proliferation of these cells. This suggest that that the differential toxicity of auranofin between the Ewing sarcoma and non-transformed cells is due to an increased reliance on active protein synthesis in the cancer cells, which is well-described in the literature (11,13,26,52–57).

Although we identified that ROS activates 4E-BP1 we expect that the mechanism of how ROS inhibits protein synthesis is complex and not limited to only activation of 4E-BP1. For example, recent work identified that ROS selectively targets and modifies cysteine residues in ribosomal proteins and, thereby, reduces protein synthesis (15). Future work will focus on delineating the contributions of these different targets of ROS and their effects on protein synthesis. In addition, we also plan to investigate the mechanism of how ROS activates 4E-BP1 in sarcoma cell lines. Work in other cell types has shown that ROS can inhibit mTOR activity and activate 4E-BP1 via activation of the peroxisome-bound tuberous sclerosis complex (TSC) and this will be a focus of future investigation (16,58,59). Finally, future work will also investigate the impact of the different types of ROS on protein synthesis.

Auranofin, an inhibitor of thioredoxin reductase, is approved for the treatment of rheumatoid arthritis (35,36). Notably, Pessetto et al. previously described in vitro and in vivo sensitivity of Ewing sarcoma cells to auranofin (23). In addition, the sensitivity of Ewing sarcoma cell lines to auranofin was dependent on the expression of the EWS-FLI1 oncoprotein, which suggests that thioredoxin reductase may target a unique vulnerability in Ewing sarcoma tumors (23). Other groups have also identified that Ewing sarcoma cells are sensitive to reactive oxygen species and to the manipulation of oxidative stress, using drugs such as fenretinide (18,21,22). Furthermore, a number of drugs commonly used in the clinic, including doxorubicin and cisplatin, are known to generate ROS, although it is currently unknown whether inhibition of protein synthesis may contribute to the mechanisms of action of these drugs (15). Recent work also identified that checkpoint kinase 1 (CHK1), which we and others previously identified as therapeutic target in Ewing sarcoma tumors, functions as a nuclear H_2_O_2_ sensor and regulates a cellular program that dampens ROS (15). Notably, CHK1 inhibition was shown to increase steady-state levels of nuclear H_2_O_2_, which suggests that combining auranofin, or other ROS-inducing drug, with a CHK1 inhibitor could enhance toxicity (15,60). These drug combinations will be investigated in future work.

Overall, our data demonstrate that reactive oxygen species activate 4E-BP1, which blocks cap-dependent protein translation, and inhibit protein synthesis in sarcoma cells. We also identified a mechanistic link between ROS and the DNA replication (RRM2) and cell cycle regulatory (PLK1) pathways.

## Author Contributions

JH designed the experiments, conducted the experiments, analyzed the data, and was a major contributor in writing the manuscript. SK and EG conducted the experiments. DG designed the experiments, analyzed the data, and was a major contributor in writing the manuscript. All authors read and approved the final manuscript.

## Acknowledgments

The authors would like to acknowledge use of the University of Iowa Flow Cytometry Facility and the DNA sequencing facility (Genomics Division of the Iowa Institute of Human Genetics).

