## Supplementary figures for "Auranofin and reactive oxygen species inhibit protein synthesis and regulate the level of the PLK1 protein in Ewing sarcoma cells"

#### Supplementary Materials

**Supplementary Figure 1.** (A) EW8 cells were treated with different doses of auranofin for 24 hours and then protein synthesis was quantified using puromycin. (B-C) EW8 (B) and TC71 (C) cells were treated with auranofin (5 μM) for six hours and then labeled with O-propargyl-puromycin for quantification of protein synthesis by flow cytometry. Cycloheximide (CHX) was used as a positive control for a drug that inhibits protein synthesis. (D-E) EW8 (D) and TC71 (E) cells were treated with hydrogen peroxide (100 μM) for three hours, auranofin (5 μM) for six hours, or auranofin (5 μM) for 24 hours. Cell viability was then quantified using 0.4% Trypan Blue (ThermoFisher), which stains dead cells, with a DeNovix Cell Drop BF automated cell counter.

**Supplementary Figure 2.** (A) BJ and RPE cells were treated with auranofin (5 μM), NAC (5 mM), or the combination for twenty-four hours. Cells were labeled with puromycin to quantify protein synthesis and then lysates were collected for immunoblotting. (B-C) Dose response curves for non-transformed (B) BJ and RPE (C) cells treated with auranofin. Cell viability was assessed 72 hours after drug was added using the AlamarBlue assay. Error bars represent the mean ± SD of three technical replicates.

**Supplementary Table 1.** Reverse phase protein array (RPPA) data for TO-4E-BP1-Ala sarcoma cell lines treated with vehicle or doxycycline, to induce expression of 4E-BP1-Ala.

**Supplementary Table 2.** Overlap of the RPPA data for proteins that are downregulated by >1.25-fold in the TO-4E-BP1-Ala sarcoma cell lines.

### Supplementary Figure 1

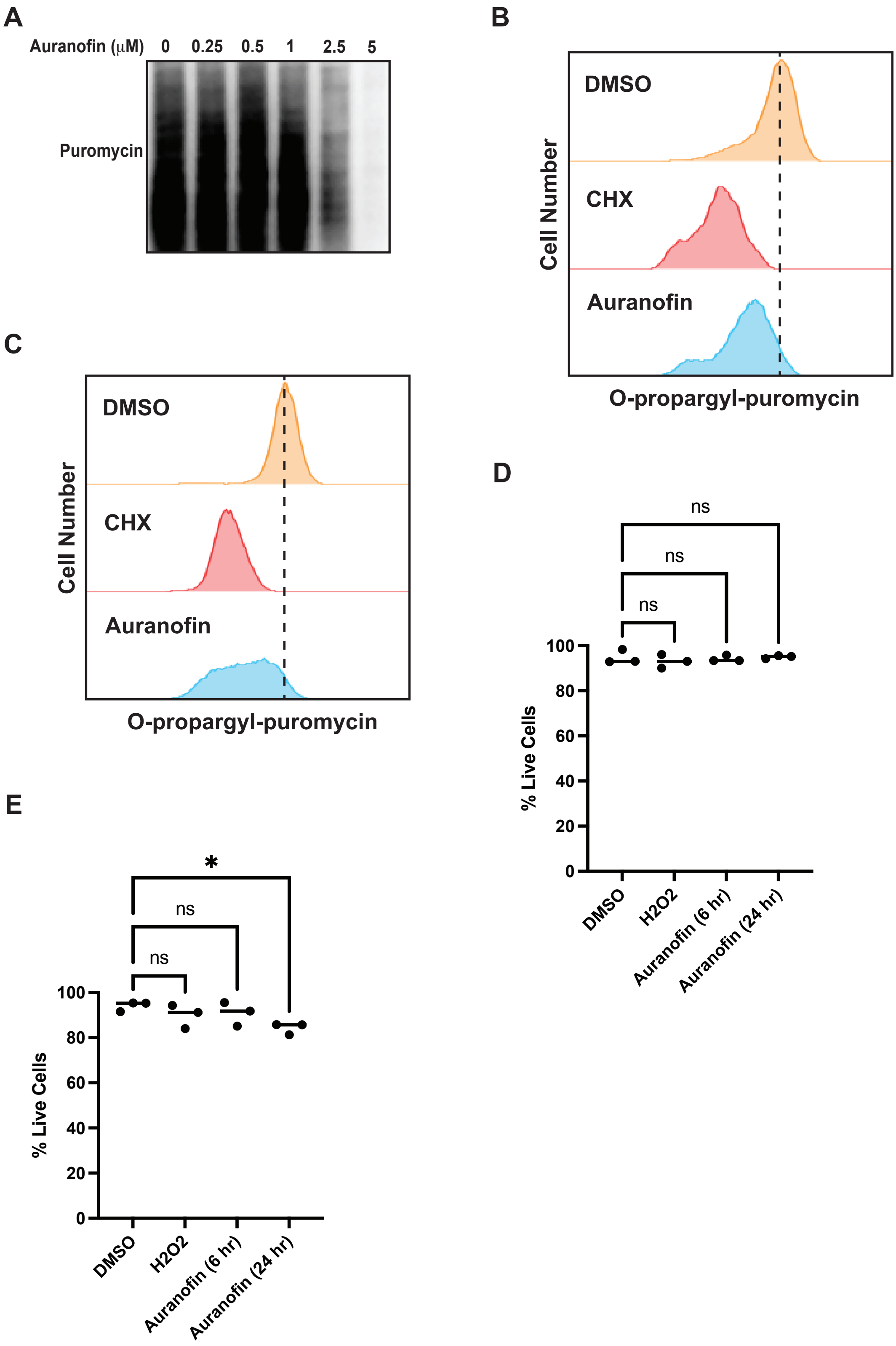

### Supplementary Figure 2

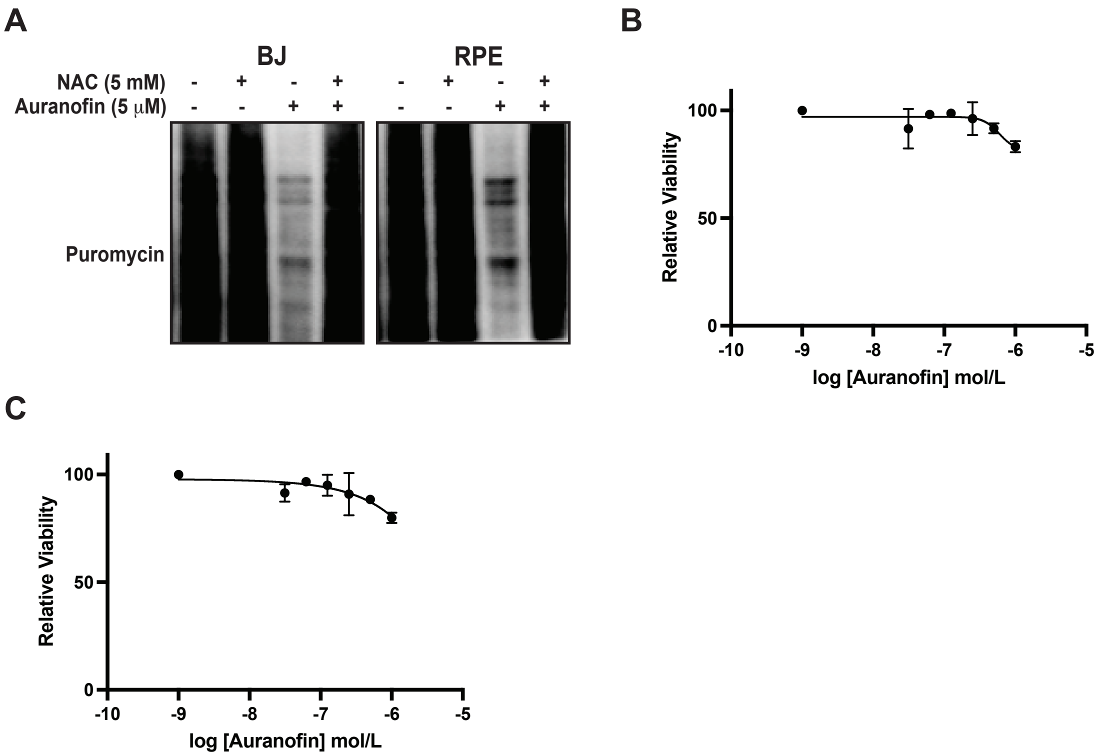
